# *Acidovorax* pan-genome reveals specific functional traits for plant beneficial and pathogenic plant-associations

**DOI:** 10.1101/2021.01.29.428753

**Authors:** Roberto Siani, Georg Stabl, Caroline Gutjahr, Michael Schloter, Viviane Radl

## Abstract

Beta-proteobacteria belonging to the genus *Acidovorax* have been described from various environments. Many strains can interact with a range of hosts, including humans and plants, forming neutral, beneficial or detrimental associations. In the frame of this study we investigated genomic properties of 52 bacterial strains of the genus *Acidovorax*, isolated from healthy roots of *Lotus japonicus*, with the intent of identifying traits important for effective plant-growth promotion. Based on single-strain inoculation bioassays with *L. japonicus*, performed in a gnotobiotic system, we could distinguish 7 robust plant-growth promoting strains from strains with no significant effects on plant-growth. We could show that the genomes of the two groups differed prominently in protein families linked to sensing and transport of organic acids, production of phytohormones, as well as resistance and production of compounds with antimicrobial properties. In a second step, we compared the genomes of the tested isolates with those of plant pathogens and free-living strains of the genus *Acidovorax* sourced from public repositories. Our pan-genomics comparison revealed features correlated with commensal and pathogenic lifestyle. We could show that commensals and pathogens differ mostly in their ability to use plant-derived lipids and in the type of secretion-systems being present. Most free-living *Acidovorax* strains did not harbour any secretion-systems. Overall, our data indicated that *Acidovorax* strains undergo extensive adaptations to their particular lifestyle by horizontal uptake of novel genetic information and loss of unnecessary genes.

## Introduction

Several soil-derived bacteria, which can colonize plant roots, directly or indirectly promote plants’ health by facilitating nutrient mobilization and uptake [1], altering plant hormone levels hormones [2] and competing with pathogens [3, 4]. At the same time, soils also harbour a large repertoire of plant pathogens. It is not entirely clear what distinguishes non-harmful root-associated commensals from pathogens, as even closely related strains have been proven to promote plant-growth or, contrariwise, to negatively affect it [5]. Several studies [6, 7] link this to loss or acquisition of genomic features [e.g., virulence factor, pathogenicity islands], resulting from mutations and horizontal gene transfer [8]. Our existing knowledge in this field is still scarce and inconsistent, as in most cases only single strains were compared, which does not allow to understand evolutionary patterns and to define a generalized model.

Bacteria of the genus *Acidovorax* belong to the group of Gram-negative beta-proteobacteria [9] and form associations with a wide range of monocotyledonous and dicotyledonous plants [10]. *Acidovorax* has been mostly considered as biotrophic pathogen [11]. However, bacteria of this genus are also known as commensal species or plant beneficial bacteria, which produce secondary metabolites and hormones promoting plant growth, as well as competing with pathogens [12]. Most common species are *A. delafieldii* and *A. facilis*, whilst the most studied pathogenic species is *A. avenae*, the agent of corn (*Zea mays*), sugar cane (*Saccharum officinarum*) and rice (*Oryza sativa*) leaf blight, orchids brown spot disease and watermelon (*Citrullus lanatus*) fruit blotch. Thus, *Acidovorax* is a valuable model organism to investigate evolutionary patterns of plant-associated bacteria and to study which genes differentiate bacteria acting as biotrophic pathogens, as commensals or plant growth-promoting bacteria [13].

In the frame of this study, we compared the genomes of isolates classified as *Acidovorax* which were obtained from roots of healthy *Lotus japonicus* ecotype Gifu plants. We tested the strains for their ability to promote plant growth of *L. japonicus* in sterilized sand and correlated differences in the genomes of the strains to the observed effects on plant growth. For a broader comparative analysis, we included additional genomes, available in public databases, belonging to strains from 15 different species of *Acidovorax* from different environments, including all major groups of plant pathogens from this genus. We studied the diversity in the pan-genome and critically evaluated enriched gene functions, secondary metabolites biosynthetic clusters in the different functional groups of *Acidovorax*.

## Material and methods

### Origin of strains and genomes

Fifty-two strains of endophytic *Acidovorax* were isolated from the root systems of healthy *Lotus japonicus* ecotype Gifu B-129 (subsequently called *L. japonicus*) grown in natural soil in Cologne, Germany. DNA from all strains was extracted and sequenced using an Illumina HiSeq 2500 device (Illumina, USA). The isolation and sequencing procedure is described in detail elsewhere [14]. Genomes assemblies are available in the European Nucleotide Archive under the accession number PRJEB37696.

In addition, 54 other genomes where obtained from NCBI. The collection includes 15 species: *A. anthurii, A. avenae, A. carolinensis, A. cattleyae, A. citrulli, A. delafieldii, A. defluvii, A. ebreus, A. facilis, A. kalamii, A. konjaci, A. monticola, A. oryzae, A. radicis, A. varianellae* and *A. spp*. According to the database, all the selected bacterial genomes originate from plant, soil and water samples. Details on the selected genomes are summarized in Supplemental Material 1. The mean level of completion for all 106 genomes was calculated to 0.986.

Based on the available metadata, we assigned the genomes to 3 behavioural phenotypes: 62 genomes belong to commensal or beneficial plant-associated strains (thereby termed “commensals”), 21 to plant pathogenic strains (thereby termed “pathogens”) and 23 to free-living strains (thereby termed “free-living”). According to the results of our *in planta* bioassays (see below), we further classified the *Lotus*-isolated strains by their ability to promote *Lotus japonicus* growth.

### Linear discriminant analysis effect sizes

We studied genomic features across the strains isolated from *L. japonicus* and correlated the obtained data with the effects detected by the *in planta* bioassays (see below). Therefore, we annotated the genomes using Anvi’o 6.2 microbial multi-omics platform [15] which computes *k-* mer frequencies and identifies coding sequences using Prodigal [16]. Afterwards, HMMER3 [17] was used to profile the genomes with hidden Markov models and to find homologous protein families in the Pfam database v33.1 [18]. Using the features occurrence frequency matrix generated by Anvi’o 6.2, we calculated the effect sizes of the features over a linear discriminant analysis (LEfSe [19]) between the groups of interest. We chose the default alpha values of 0.05 for the factorial Kruskal-Wallis test (KW) and the pairwise Wilcoxon test (W) and a threshold of 2.0 for the Linear Discriminant Analysis (LDA) logarithmic scores.

### Pan-genome analysis

We used Anvi’o 6.2 further to pre-process the genomes and reconstruct *Acidovorax* pangenome [20]. We inferred the pan-genome with the following options: DIAMOND in sensitive mode to calculate amino-acids sequences similarity, Minbit parameter to 0.5, minimum occurrence of gene clusters to 2, MCL inflation parameter to 7. We separated gene clusters into occurrence frequency classes. Kernel gene clusters were defined as present in 106 genomes. Core gene clusters were defined as present in 100 to 105 genomes. Shell gene clusters were defined as present in 16 to 99 genomes. Cloud gene clusters were defined as present in 2 to 15 genomes. Singleton gene clusters were defined as present in only 1 genome and were excluded from the downstream processing to reduce computational load. We calculated functional and geometrical homogeneity for each gene cluster and genome average nucleotide identity using pyANI [21]. We calculated a functions occurrence frequency matrix and functions enrichment scores for gene clusters’ functions by assigning each gene cluster to the closest Pfam.

### Comparative genomics

We performed all downstream analysis in R (v4.02, code available on https://github.com/rsiani/AVX_PGC). We calculated total length, GC%, number of genes, gene clusters and singleton gene clusters, average gene length and number of genes per kb (gene density) for each of the genomes. We assessed the significance of differences by Pairwise Student’s t-test with Holm-Bonferroni’s correction for multiple testing (*p* < 0.05).

We converted the function frequency occurrence matrix (3036 Pfams) to a binary presence/absence matrix and filtered zero variance (757 Pfams), near zero variance (1668 Pfams) and correlated variables (0 Pfams) to remove redundancy (resulting in 1368 Pfams). We performed a Principal Component Analysis (PCA, R package: “FactoMineR”, v2.3) to ordinate the strains based on their functional similarity and tested the significance of the clustering by PERMANOVA (package: “vegan”, v2.5-7). We isolated the features responsible for most of the variance explained by extracting the highest contributors for each of the first 3 components.

From the enriched feature profiles predicted by Anvi’o 6,2, we manually curated a list of features putatively correlated with plant-association. Enrichment score and adjusted significance value were used to assess the relevance of the features across the groups. We retrieved secondary metabolites biosynthesis gene clusters using AntiSMASH 5.0 [22]. We launched a local instance of the program with no extra features at a “relaxed” strictness level, running only the core detection modules. At this strictness level, well-defined clusters and partial clusters are detected and annotated against the antiSMASH database.

We built a classification model using neural networks with feature extraction (R package “caret”, v6.0-86), a model employing a preliminary feature extraction step to reduce redundancy, while preserving information and decreasing computational load [23]. We trained the model over 90 % of the strains with “leave-one-out” cross-validation over 1000 repeats, using accuracy scores to select the optimal parameters. The fit of the model was evaluated by predicting labels for the remaining 10 % of the strains and again against the whole dataset Results were assessed by calculating confusions matrix for both predictions.

### *In planta* bioassays

We prepared seeds of *L. japonicus* by sandpaper scarification and surface sterilization with a 10% DanKlorix Original (CP GABA GmbH, Germany) and 0.1% sodium dodecyl sulfate (SDS) solution. We germinated the seeds on a 0,8 % water-agar plate for 3 days at 22 °C in the dark followed by 4 days at 22 °C in the light at 210 μM/m^2^s intensity (growth cabinet PK 520-LED, Polyklima).

We transferred plantlets to pots (Göttinger 7 x 7 x 8 cm, Hermann Meyer KG, Germany) filled with washed and autoclave sterilized quartz-sand (Casafino Quarzsand, fire-dried, 0,7 – 1,2 mm, BayWa AG, Germany) at a density of 5 plants per pot. Pots were supplied with a thin layer of synthetic cotton at the bottom to prevent the sand running out through the holes. 44 of the *Acidovorax* strains isolated from healthy *L. japonicus* plants were individually pregrown in 50 % TSB media at 30°C overnight and diluted to an OD_600_ of 0,001 using 1/3 BNS-AM medium (Basal Nutrient Solution [24] modified for arbuscular mycorrhiza (0.025 mM KH2PO4)). 25 ml of this bacterial suspension was applied to the pots. Finally, 20 ml of the sterile 1/3 BNS-AM-bacterial solution was added. Plants were grown at 60 % air humidity and light intensity of 150 μM/m^2^s. The photo-period was set to 16 h light/ 8 h dark long day with a temperature cycle of 24/22 °C. For each bacterial strain tested, four replicate pots were prepared.

After 9 weeks of growth, we harvested the plants and determined root and shoot length, wet weight and dry weight after freeze drying (Alpha 1-2, Martin Christ Gefriertrocknungsanlagen GmbH, Germany). Statistical significance was assessed by Pairwise Student’s *t*-test adjusted for multiple comparisons using the Holm-Bonferroni method (*p* < 0.05).

## Results

### *Acidovorax* strains isolated from healthy *L. japonicus* roots diversely affect their host

We compared growth metrics of *L. japonicus* inoculated with different strains to evaluate the outcomes of the plant-association. Overall, 34 out of 44 tested strains showed significant effects (*p* < 0.05; Figure 1) when considering all the collected measurements of plants’ growth (n = 25). Twenty-one strains displayed a positive effect on at least one metric. Nine strains displayed a negative effect on plant growth, markedly on fresh weight measurements. Four strains displayed contrasting effects on different metrics and 10 had no significant effect on growth. From the 21 strains that resulted in improved plant-growth, 7 strains displayed a more consistent effect across all the replicates on at least one of the considered metrics (*p* < 0.001). We selected these robust growth-promoters for a follow-up genomics comparison.

**Fig. 1:**
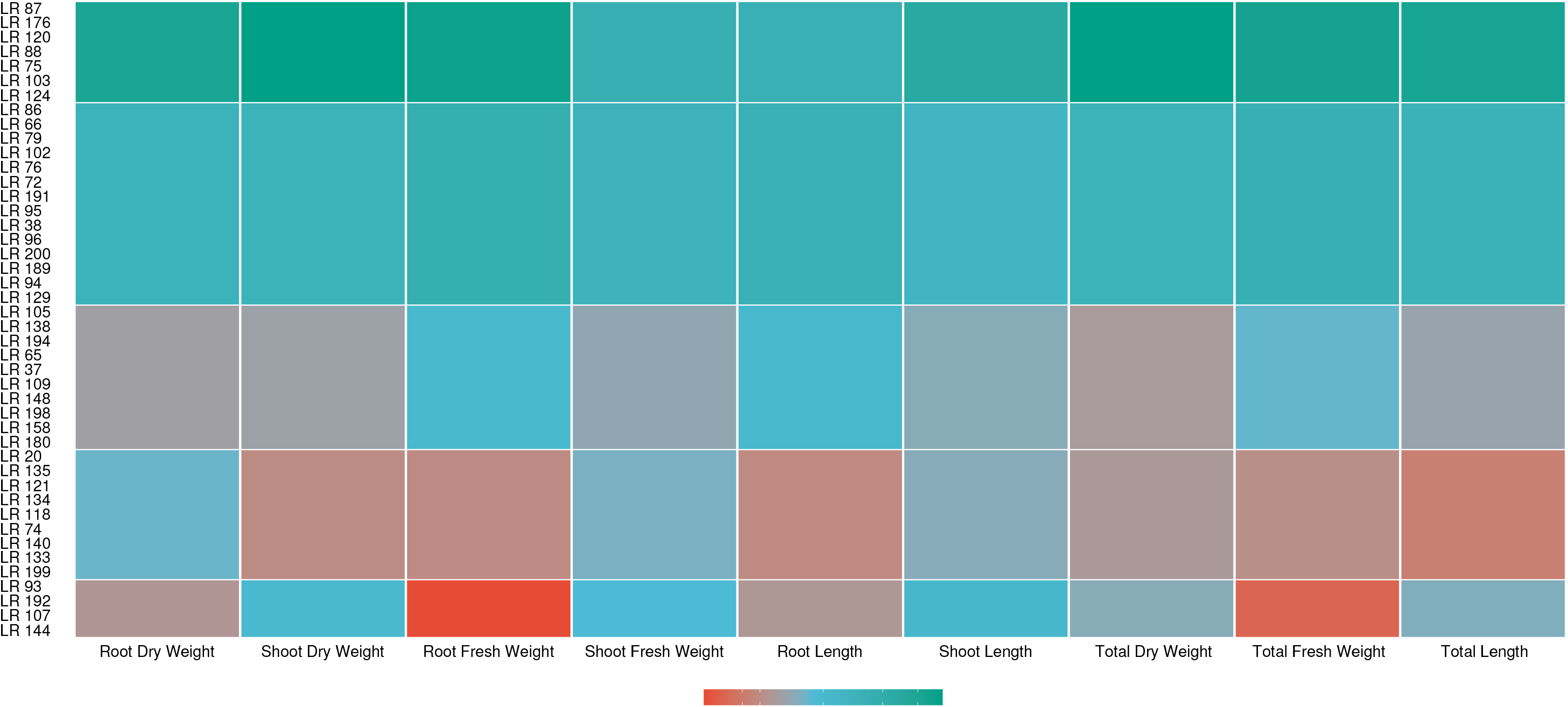
Heat-map showing the effects of single-strains on *Lotus japonicus* growth parameters as observed in the *L. japonicus* growth bioassays. Measures are shown as the median, for each observed group, of the percentage change from the control.

### Nineteen Pfams discriminate the robust plant growth-promoting *Acidovorax* strains

To isolate genomic differences correlated with the outcomes of the *in planta* bioassays, we calculated linear discriminant analysis effect sizes on the function occurrence frequency table and identified 19 Pfams discriminating the 7 robust growth-promoters from the remainder of the *Lotus*-isolated strains (KW *p* < 0.05, W *p* < 0.05, logarithmic LDA score > 2.0, Figure 2; Table 1). Out of those, 15 Pfams were enriched in the genomes of robust growth-promoters, related to 4 broad functional categories: chemotaxis by sensing and uptake of organic acids and sugars (tripartite tricarboxylate transporters, TTTs, and periplasmic binding proteins, PBPs), colonization through synthesis and detoxification of plant secondary metabolites (aldolase, aldoketo reductase, rhodanase, amidase and mannosyltransferase), competition by synthesis, secretion and detoxification of antimicrobial compounds (beta-lactamase, polyketide cyclase, tetrapyrrole corrin-porphyrin methlyase, dynein-related domain and dienelactone hydrolase) and transcriptional regulation (FCD domain, IclR domain and sigma 70 factor). We found 4 Pfams which were less abundant in the genomes of robust growth-promoters, also implicated in chemotaxis (Tripartite ATP-independent periplasmic transporters, TRAPs), colonization (FliO and the FG-GAP repeat) and regulation (metallo-peptidase M90).

**Fig. 2:**
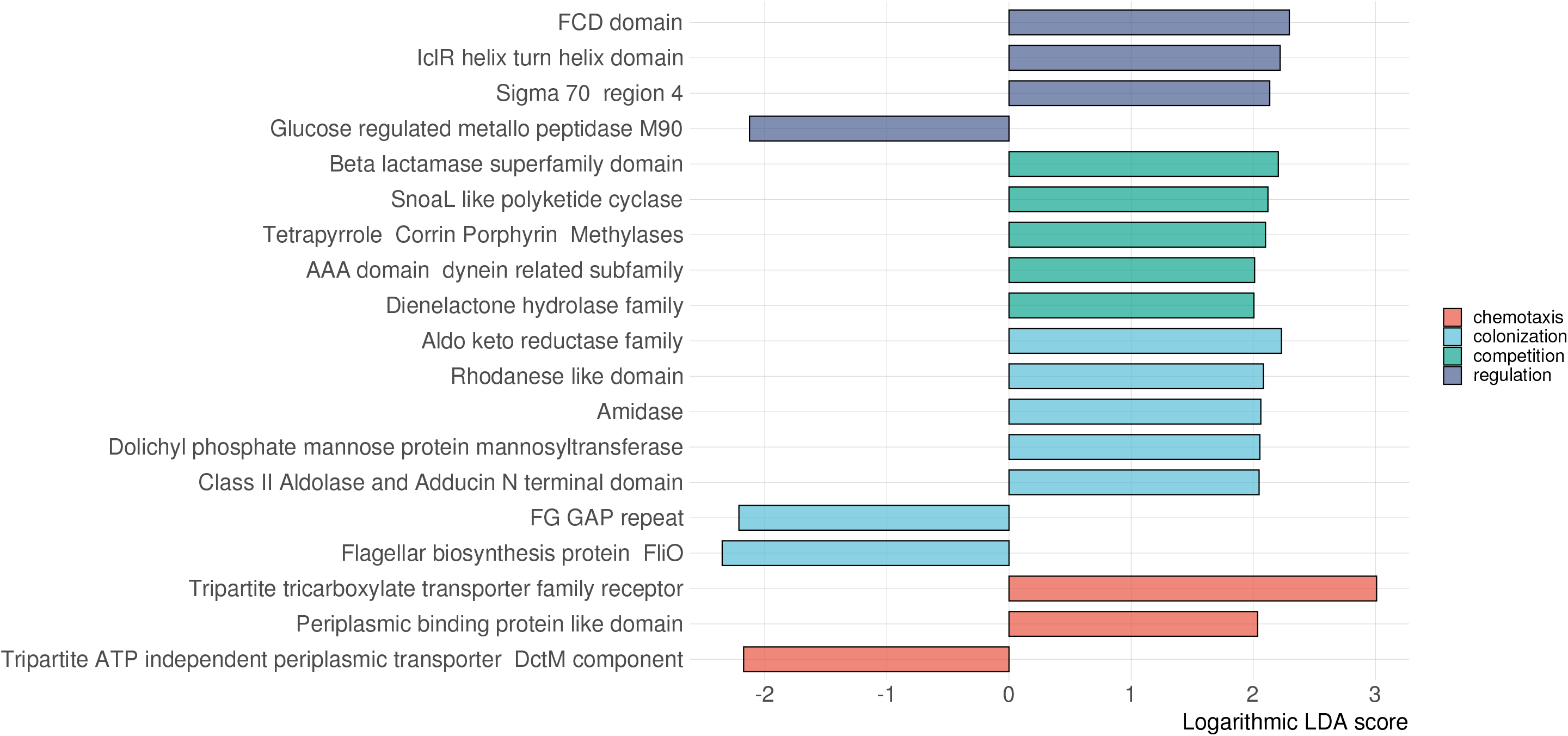
Bar-plot of the logarithmic scores from the Linear Discriminant analysis for the discriminant Pfams. Pfams are assigned to 4, color-coded, functional groups.

### *Acidovorax* has an open pan-genome

We reconstructed the *Acidovorax* pan-genome from 106 genomes (Figure 3), representing 15 out of the 23 classified *Acidovorax* species [51], along with several yet unclassified strains, and capturing a large share of the genomic variability in the genus. The pan-genome contains a total of 523555 genes grouped in 34146 gene clusters, of which 2 % belonged to the kernel (present in all of the genomes), 3% belonged to the core (present in 100 to 105 genomes), 13% belonged to the shell (present in 16 to 99 of the genomes), 45 % belonged to the cloud (present in 2 to 15 genomes) and 37 % were singleton (present in one genome only). Ninety-five percent of the gene clusters were classified as accessory, while 5 % are shared by all or most of the genomes. By fitting our results to a Heaps’ Power-law regression model [52] we obtained a γ < 1 of 0.48, which defines *Acidovorax* pan-genome as “open”, with positive rates of novel genes discovery for every additional genome included.

**Fig. 3:**
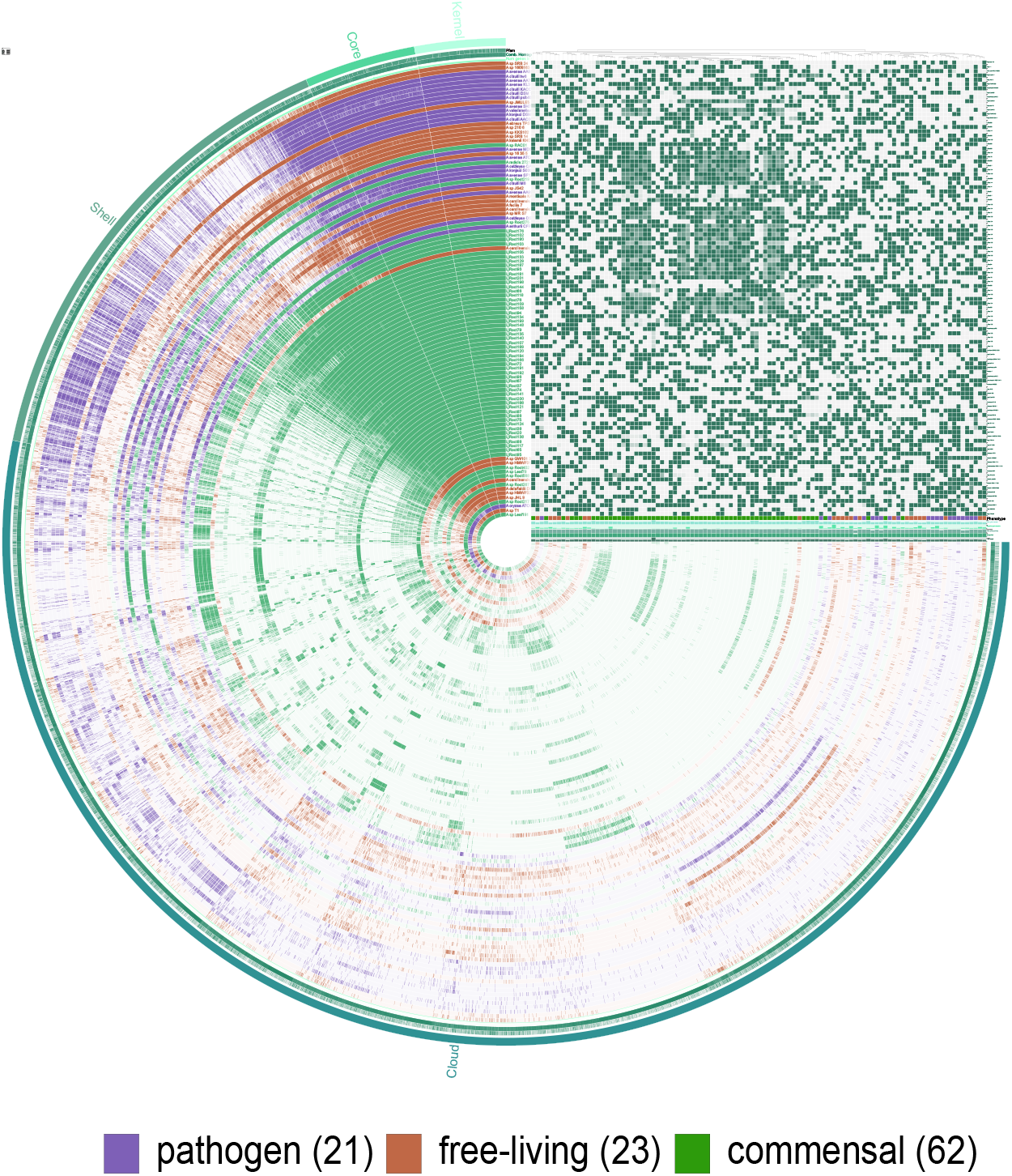
Anvi’o 6.2 circular representation of *Acidovorax* pan-genome and heat-map of average nucleotide identity across the genomes.

### Pathogenic *Acidovorax* strains have longer genomes but less genes

We calculated total length, GC%, number of genes, gene clusters and singleton gene clusters, average gene length and number of genes per kb (gene density) for each of the genomes (Figure 4a, Supplemental Material 2). On average pathogens had longer genomes (5.38 mb, SD = 0.27) than commensals (5.21 mb, SD = 1.03, *p* < 0.01) and free-living (4.72 mb, SD = 0.53, *p* < 0.0001, Figure 4b). However, when testing for differences in gene density, pathogens had less genes per kb (0.86, SD = 0.09) than both commensals (1.01, SD = 0.05, *p* < 0.0001) and free-living (0.94, SD = 0.03, *p* < 0.001, Figure 4c).

**Fig. 4:**
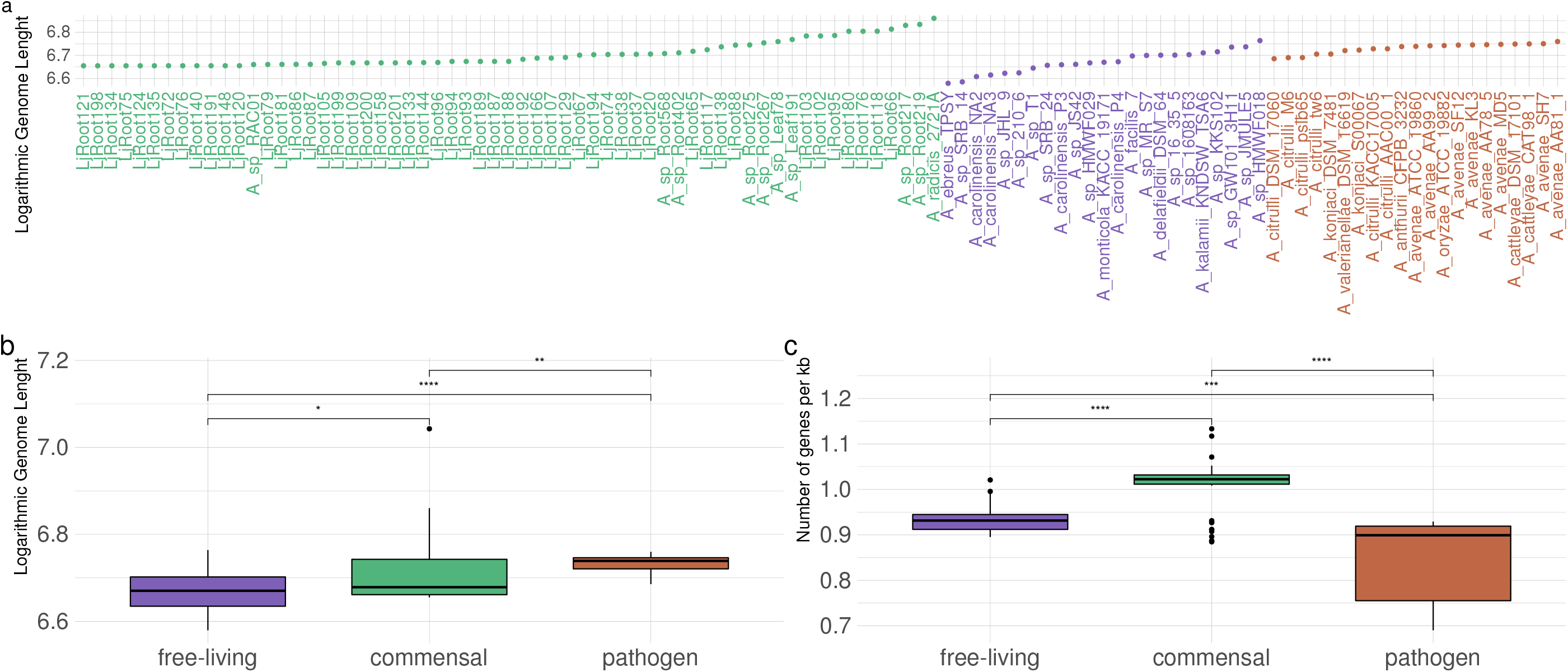
a) Dot-chart of individuals logarithmic genome length. b) box-plot of groups logarithmic genome length. c) box-plot of groups gene density expressed as number of genes per 1000 base-pairs. Significance was assessed by Student’s t-test with Holm’s correction (*p* < 0.05).

### Strains separate according to behavioural phenotype

We performed a PCA on the function presence/absence matrix and found that the first 3 dimensions explained 47 % of the variance in the matrix (24.3 %, 12.4 % and 10.3 %). The individual strains clustered according to their observed behavioural phenotype (*p* < 0.05), with pathogens grouping in a neatly separated cluster and a partial overlap between free-living and commensal strains (Figure 5ab). Notably, no clustering was visible when considering the original habitat of the strains. We extracted the highest contributing variables for the first 3 components (Figure 5c, Table 2).

**Fig. 5:**
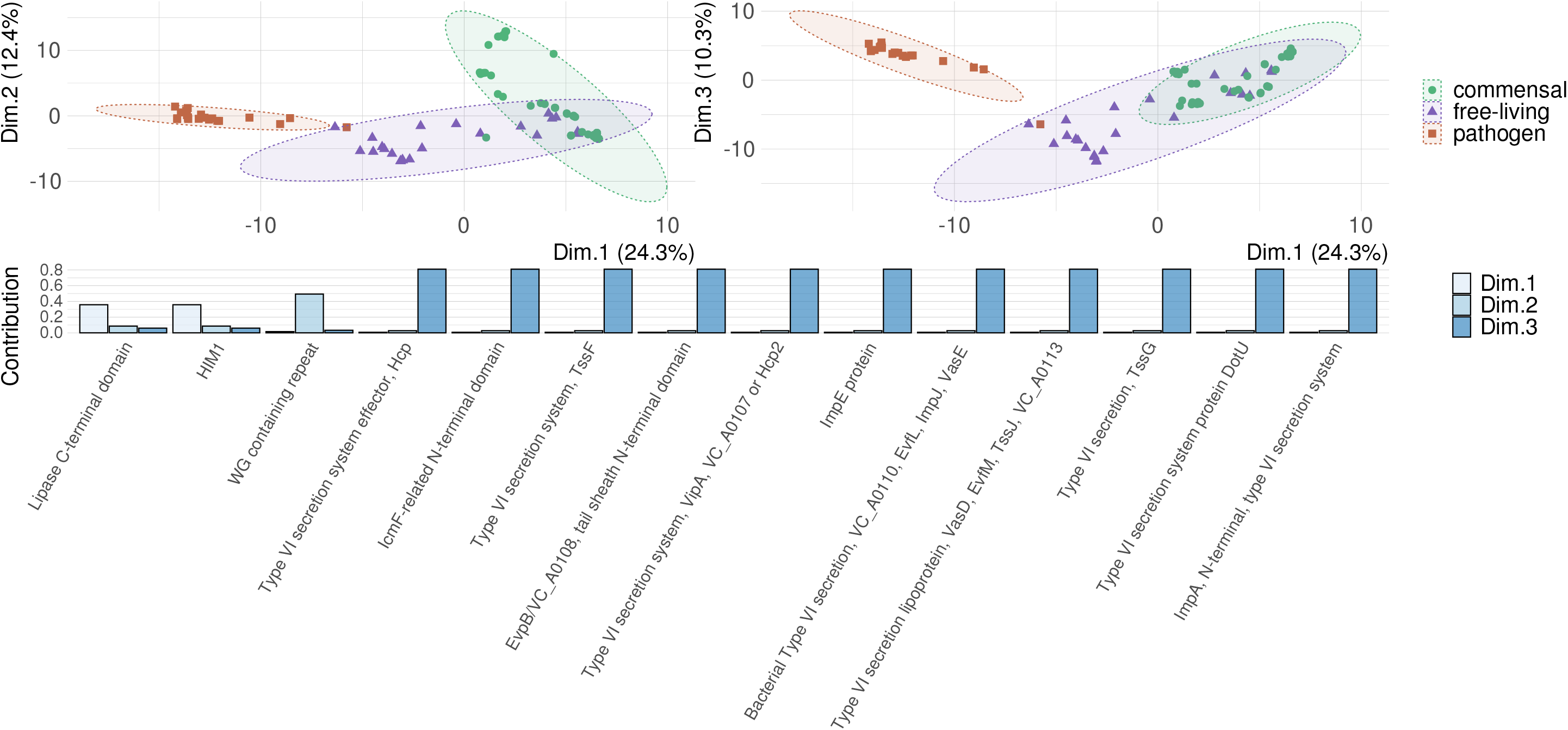
a) Ordination plot for the first and second component of the principal component analysis (PCA) b) Ordination for the first and third component of the PCA. c) Bar-plot of the Pfams with the highest contribution for first, second and third component of the PCA.

This approach retrieved a total of 14 Pfams displaying a strong variance across the groups. Two of these, the C-terminal domain of a secreted lipases and the mutagenesis inducer HIM1, contributed the most to the first component and were found in all of the commensals, 35 % of the free-living and none of the pathogens. A third Pfam, characterized by a WG repeat motif, was only retrieved in 40 % of the commensals and accounted for a high proportion of the second component explanatory power. Finally, the highest contributor to the third component was a putative genomic island, encoding 11 elements of the type VI secretion-system, present in 95 % of the pathogens, 61 % of the commensals and only 13 % of the free-living strains.

### Pathogenic and commensal *Acidovorax* strains have a distinctive set of features

We performed a function enrichment analysis to isolate features unique or more represented in the different groups (Supplemental Material 3). 1243 out of 3036 Pfams were significantly enriched in 1 or 2 groups (adjusted *q* < 0.05). We further investigated enriched functions from plant-associated strains only: 371 Pfams strongly associated with commensals (*q* < 0.05) and 303 Pfams strongly associated with pathogens (*q* < 0.05) (Figure 6a). From these, we manually curated a list of 54 features (Table 3) putatively involved in plant-microbe and microbe-microbe interactions (Figure 6b). We assigned them to 4 families based on known performed function: 22 effectors, 13 hydrolytic enzymes, 12 motility functions and 7 secondary metabolites.

**Fig. 6:**
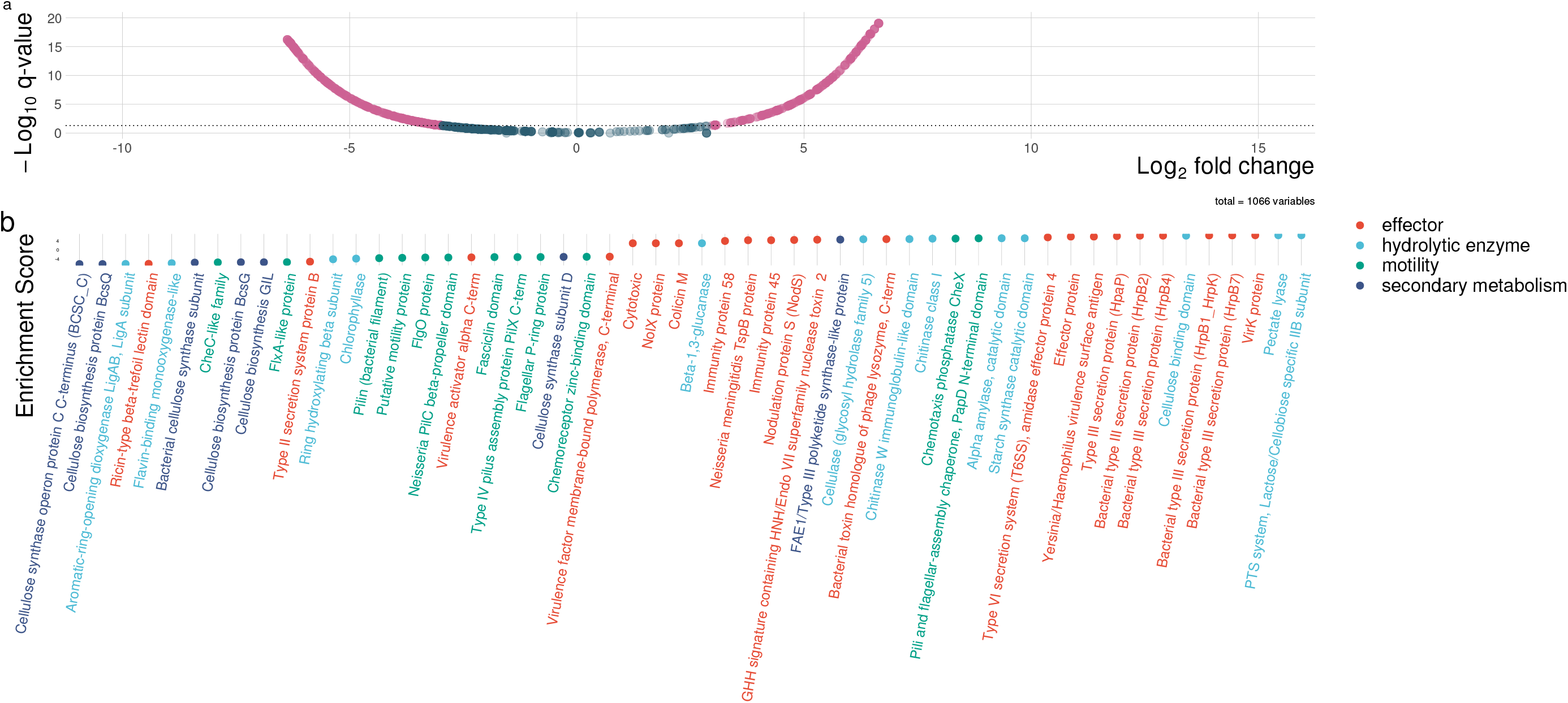
a) Volcano plot displaying Pfams enriched in commensal (left) and pathogenic strains (right). On the x axis is represented logarithmic fold change and on the y axis adjusted *q*-value. Dots are represented in red when crossing the significance threshold of 0.05. b) Dot-plot of the curated list of enriched Pfams related to host-microbe or microbe-microbe interaction. Each dot is color coded based on the functional family of assignment.

Eighteen effectors and known virulence factors were enriched in pathogens, among which, notably, members of the HrpB gene of the type III secretion system, and its regulatory element HpaP. Four effectors were also identified in commensals, including a type II secretion-system protein.

Among the hydrolytic enzymes, 9 different Pfams were found to be enriched in pathogens, all related to plant and fungal polysaccharides degradation, including components of pectate lyases, amylases, chitinases, cellulases and glucanases. Four hydrolytic enzymes Pfams were enriched in commensals, but mainly responsible for the degradation of complex bioactive compounds, such as flavonoids, chlorophyll and aromatic-ring backbones.

Interestingly, 10 Pfams related to diverse motility structures, namely flagella and pili, and chemotaxis were predominantly found in commensals. For comparison, just 2 motility related Pfams were enriched in pathogens.

Among those enriched in commensals, we also found 6 Pfams related to bacterial cellulose biosynthesis and 1 polyketide-synthetase Pfams enriched in pathogens.

Using AntiSMASH 5.0, we retrieved a repertoire of 15 biosynthesis gene clusters (BGCs) among the bacterial genomes (Figure 7). The most commonly retrieved BGCs were those for bacteriocin and terpene production. Interestingly, phosphonate and all non-ribosomal peptidesynthetase BGCs (NRPS, NRPS-like, NRPS-T1PKS) were almost unique to pathogens and retrieved in more than half of the pathogenic strains.

**Fig. 7:**
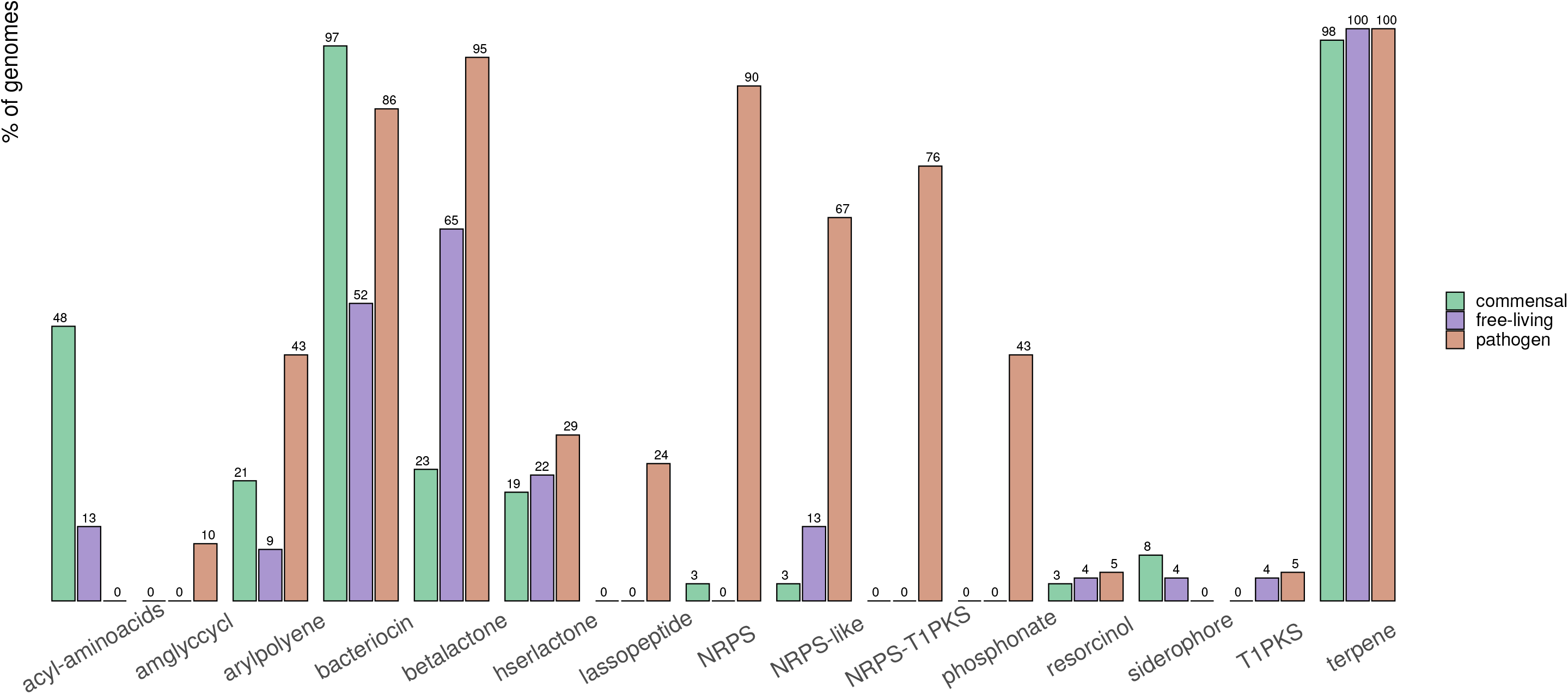
Bar-plot of the percentage of genomes per group featuring each of the retrieved biosynthesis gene clusters.

### Genotypes can predict the behaviour of the strains

We tested whether a model could predict a strain behavioural phenotype from its gene repertoire. We trained a neural network with feature extraction classifier on the gene function’s presence/absence profiles of 90% of our strains (n = 96). The optimal model, which we selected according to accuracy value, registered a mean balanced accuracy = 0.93 and an area under the curve (AUC) = 0.97, with the selected final values of size = 3 and decay = 0.1. We tested the predictive power of the model against the remainder 10% of the strains (n = 10) and registered accuracy = 1, sensitivity = 1, specificity = 1, kappa = 1, 95% confidence interval = 0.69 to 1 and *p* = 0.006. When testing against the whole collection (n = 106), all metrics remained unchanged but for the 95% confidence interval = 0.97 to 1 and *p* = 2.2e-16.

## Discussion

### Robust growth-promoting strains share traits which allow a better use of phytochemicals

Our *in planta* bioassays showed a wide range of effects associated with the presence of diverse *Acidovorax* isolates obtained from healthy roots of *L. japonicus*. We compared the genomes of 7 robust growth-promoters against the remainder of the isolates and retrieved 19 discriminant Pfams related to different aspects of plant-microbe and microbe-microbe interactions: chemotaxis towards root exudates, metabolism of plant secondary metabolites, antagonistic competition and transcriptional regulation.

Organic acids are a major component of root exudates [110]. Microbial transporters, such as ABC-type transport systems, TRAPs and TTTs, are essential to make use of these carbon rich compounds and a number of studies confirmed that their presence is correlated to improved colonization in both plant-growth promoters [111] and pathogens [32]. Many transporters are highly specific for individual substrates. For example, TTTs, which are enriched in robust-growth promoters in our study, show a higher affinity towards citrate compared to TRAPs. As shown by metabolic profiling [112], citrate is more abundant than other organic acids in *L. japonicus* root tissue and improved sensing and uptake of citrate could translate in an evolutionary advantage of microbiota towards the colonization of this plant. Our robust growth-promoters also shared an enrichment of several features related to the metabolism of various plant-derived compounds, such as benzoic acids, carbonyls, cyanide and rare sugars. Some of these traits have been already correlated to improved plant-association in taxonomically diverse bacteria [13, 113, 114]. Furthermore, we observed the enrichment of an amidase involved in the biosynthesis of indole-3-acetic acid and of a known regulator of GABA uptake [115]. The role of auxin in plant-growth promotion by bacteria has been thoroughly characterized [116] and GABA has been more recently discovered to act as signal from plants to their associated microbiota [117, 118]. Finally, we found enriched components of diverse regulatory pathways and of both antibiotic synthesis and degradation, which could help the robust growth-promoters to not only better colonize the plant by degrading plant-derived phytochemicals with antimicrobial properties, but also to gain competitive advantage over other microbial populations.

Overall, we found evidence of genomic differences among robust growth-promoting bacteria and the remainder of *Acidovorax* isolates, suggesting improved chemotaxis, competitive traits and interaction with the plant metabolism and hormonal balance, possibly leading to a stronger association between host and colonizers and explaining the better growth outcomes observed *in planta*.

### The evolutionary trajectory towards pathogenic plant-association

*Acidovorax* strains are naturally found to occupy widely diverse niches. In the frame of this study, we reconstructed their pan-genome to explore the importance of genomic features across the whole behavioural spectrum. Pan-genome analyses have become a staple to estimate the complete gene repertoire accessible to an organism and to understand genotype variations and evolution in a broader context [119, 120]. We found that the *Acidovorax* pangenome is largely made up by accessory or unique gene clusters (95 %) and predisposed to a continuous inflation, a predictive feature of functionally flexible organisms [121], with each occupied niche being a potential source of novel genomic traits.

Specialization has been associated with smaller genomes and previously reported in pathogenic strains [121, 122]. Therefore, we studied *Acidovorax* genome size and density, expecting to observe evidence of reductive evolution. For example, Merhej and colleagues analysed a comprehensive collection of 317 bacterial genomes to trace the evolutionary path from free-living to specialized intracellular strains [123], uncovering “massive gene loss” as a consequence of more gene loss than horizontal gene transfer (HGT) events. Mainly in isolated niches like the interior of roots, HGT rates are considered as low due to the missing interaction with other microbiota. Surprisingly, in our study, the genomes of pathogenic *Acidovorax* were the largest, yet showed a significantly lower gene density than both free-living and commensals strains. Similar data was also observed for *Rickettsia prowazekii* [124], which features more non-coding sequences than any of its close, non-pathogenic, relatives. In their review on pathogenomics, Georgiades and Raoult argued that true bacterial speciation is only observed in the contest of segregated niche, such in the case of obligated parasitism [125]. For Acidovorax, Fegan [10] listed 28 known Graminaceae hosts for *Acidovorax avenae* subsp. *avenae* and 10 Cucurbitaceae natural hosts and 2 Solenaceae for *Acidovorax avenae* subsp. *citrulli*. Thus, *Acidovorax* pathogens seems still far from true speciation, as suggested by their wide choice of hosts, and, likely, have already endured evolutionary driven gene losses, whereas extensive genome reductions still have to occur.

### Major functional differences across the groups

Principal component analysis revealed that pathogens differ in their genomes from commensal and free-living strains. Among the differences that contributed the most to the separation, we found that the lipase C-terminal domain and the mutagenesis inducer HIM1 were both enriched in commensal strains. Assis and colleagues investigated the phylogeny of lipases in bacteria and found orthologous of the same secreted lipase ubiquitous among plant-associated strains, including *Acidovorax* [126]. Pathogens may be less dependent on lipases, as, during infection, plant-derived lipids may be of minor value to them, as they preferentially hydrolyse carbohydrates to cover their nutritional needs.

The presence of type VI secretion-systems has been strongly correlated with plant-association in Gram-negative bacteria [127] and has been in observed in pathogenic and commensal strains alike. Similarly, in *Acidovorax*, we retrieved a putative type VI secretion-system genomic island in both groups of plant-associated bacteria (commensals and pathogens, but only in 13 % of the free-living strains). For the pathogens, we also identified an operon encoding components of the type III secretion-system, absent from the other groups.

The distribution of these Pfams across the groups suggests the recruitment of commensals from the larger pool of free-living strains through horizontal gene transfer of a single genomic island encoding elements of the type VI secretion-system and further specialization of commensals into pathogenic strains, through gene losses and acquisition of type III secretion-systems.

We observed in the commensals an enrichment of several motility-associated domains. Pallen and Wren [128] postulated the loss of motility as a common side-effect of intracellular endosymbiosis, a phenomena recently witnessed in several emerging pathogens. It stands to be clarified whether this adaptation occurs for disuse, to prevent recognition by the host immune system or both. For commensals however, which are depending on chemotaxis to make use of the plant-derived phytochemicals, motility is an essential function.

Finally, pathogens showed an enrichment of polyketide synthases and non-ribosomal peptide synthases, which have been both extensively researched due to their promising role in pharmaceutical applications for the production of diverse antimicrobial, immunosuppressive and cytostatic compounds [129, 130]. Pathogenic *Acidovorax*, thus, have access to a wider array of secondary metabolites, likely necessary for microbe-microbe competition [131] and possibly to interact with the host.

### Conclusions

The focus of this research study has been to assess the role of genomic traits across the *Acidovorax* behavioural spectrum and to evaluate which genomic features can discriminate plant-pathogenic from commensal and plant-growth promoting strains. We have reported many discriminant traits through association of plant growth data with *in silico* pan-genome-wide comparisons. The robustness of our analysis was also supported by our neural network classifier, which accurately matched each individual to its observed phenotype by using the information encoded in the pan-genome, validating our hypothesis. Researchers in the field of genomics and microbiology have been investigating the potential of machine learning applications to extract meaningful patterns from the high-dimensional data generated by Next-Generation Sequencing [132, 133, 134, 135]. Even though our method was tested on a single genus, it shows promises for generalization and the advancement of genotype-based, high-throughput phenotyping and monitoring of emerging pathogens.

However, our data is based on genomic predictions and the importance of the described Pfams, both for plant-growth promoting commensals as well as plant-pathogens, needs to be verified firstly by analysing bacterial transcriptomes during the interaction with the plant, followed by targeted mutagenesis and analysis of the impact of bacterial mutants on survival in the rhizosphere and plant growth.

## Authors statement

The authors declare no financial conflicts of interest.

## Acknowledgement

Michael Schloter and Caroline Gutjahr were supported by grants from the Deutsche Forschungsgemeinschaft [DFG; SCHL446/38-1 and GU1423/3-1] respectively the frame of the SPP2125 “Deconstruction and reconstruction of the plant microbiota [DECRyPT]”. We thank Ruben Garrido-Oter for making the isolates available for the *L. japonicus* bioassays and the respective sequenced genomes for the genomics comparison We highly appreciate the discussions with Martin Parniske and Corinna Dawid. We thank M.H.H. Nguyen for help with the *L. japonicus* bioassays.

